# Molecular characterization of the heterozygous loss of function mutations in the X-linked PCDH19 gene causing PCDH19-Cluster Epilepsy

**DOI:** 10.64898/2026.03.16.712128

**Authors:** Shivang Khandelwal, Ela Elyada, Reut Japha, Manar Abu Diab, Anurag Prabhu, Zahava Siegfried, Rotem Karni

## Abstract

PCDH19-Cluster Epilepsy (PCDH19-CE) is a rare neurological disorder caused by mutations in the *PCDH19* (Protocadherin-19) gene and is characterized by early-onset seizures and cognitive impairment. In contrast to most X-linked disorders, *PCDH19* mutations predominantly affect heterozygous females, while hemizygous males are largely spared. Although advances have been made to understand the pathological mechanism underlying PCDH19-CE, key downstream targets and compensatory pathways are yet to be elucidated. Using CRISPR/Cas9 technology, we generated both a mouse model of PCDH19-CE and a human embryonic stem cell (ESC) model. Transcriptomic analysis identified genes that were differentially expressed in the brains of heterozygous (*Pcdh19*^WT/mut^) female mice compared with wildtype (WT) and homozygous (*Pcdh19*^mut/mut^) female mice. Pathway analysis of these differentially expressed genes (DEGs) revealed enrichment in pathways involved in neuronal development, ion channel activity, synaptic development and neuronal signalling. Neurons differentiated from human ESCs carrying a *PCDH19* mutation exhibited similar gene expression patterns, with heterozygous neurons displaying a distinct expression pattern compared to both WT and homozygous mutant neurons. In contrast to the molecular phenotype, neurons derived from homozygous mutant cells showed highly elongated neurites while neurons from heterozygous cells showed intermediate neurite elongation. This suggests that neurite morphology correlates directly with levels of WT PCDH19. Overall, our findings indicate that heterozygous *PCDH19* mutations are associated with defects in the expression of genes involved in developmental, signalling, and neuronal pathways in both mouse and human disease models, while certain morphological phenotypes appear to depend on the levels of WT PCDH19.

## Introduction

PCDH19-Cluster Epilepsy (PCDH19-CE) is a rare genetic disorder responsible for causing severe epileptic seizures. It is the second most clinically relevant genetic disorder causing epilepsy after Dravet Syndrome, which is caused by a mutation in the *SCN1A* gene^1,2^. Genetic mutations in the *PCDH19* gene lead to loss of expression, or the formation of a defective protein, which is responsible for PCDH19-CE. PCDH19-CE patients suffer from neurocognitive and neuropsychiatric disabilities, epileptic seizures, autism spectrum characteristics and other comorbidities^3^. PCDH19-CE manifests symptoms mainly in heterozygous females and spares hemizygous males^4^. In females with a *PCDH19* mutation on one of the alleles, random X-inactivation leads to the formation of two types of cell populations: One that is wildtype (WT) for *PCDH19*, and the other that is mutant for *PCDH19*. In neurons, these two different cellular populations have problems forming synapses with each other, thereby leading to impaired synaptic transmission and ultimately epileptic seizures^3^. This specific model is termed “cellular interference” where there is impaired communication between two populations of cells^4,5^. Although hemizygous males are spared, there are some cases where males are affected due to the presence of a somatic mutations and mosaic pattern of expression^6^.

The *PCDH19* gene is located at locus Xq22.1 and contains six exons. The first exon encodes for more than half of the protein, including integral regions such as the six extracellular and transmembrane domains. The PCDH19 protein is 1148 amino acids long and belongs to the delta-2 protocadherin subclass of the cadherin superfamily and is extensively expressed in the brain^7,8^. PCDH19 is a transmembrane protein which is localised in neurons and actively plays a role in calcium dependent cell-cell adhesion^3^. In addition to its role in cell-cell adhesion, PCDH19 is known to be involved in critical neurodevelopmental processes including cell migration, axonal growth, signal transmission, and synaptogenesis^9,10^. Recently, PCDH19 was shown to act as a transcription factor for regulating the expression immediate early genes after undergoing NMDA-dependent proteolytic cleavage^11^.

In the past few years, advancements have been made in understanding the basic molecular mechanism of PCDH19-CE. Mouse, rat, zebrafish, as well as xenopus models have been developed in an attempt to study PCDH19-CE^3,8,12–20^. These animal models have added to our understanding of the mechanism of PCDH19-CE. In contrast to the cellular interference model, Xenopus studies have revealed that both heterozygous mutant *PCDH19* animals or complete *PCDH19* knockdown animals exhibit seizure-like behaviors^16,21^. Further, a zebrafish model showed reduced numbers of inhibitory interneurons leading to hyperexcitability in both, mosaic and non-mosaic models, also contrary to the previously described theory of cellular interference as the cause of PCDH19-CE^17,22^. A mosaic rat model generated using in-utero electroporation of CRISPR exhibited a human-like phenotype, revealing the role of PCDH19 signalling in the hippocampus and cortex in neuronal migration, heat-induced epileptic seizures, and cognitive impairments^18,23^. Although mice models for PCDH19-CE do show differences at the molecular level, they do not fully recapitulate the seizures and behavioural deficits exhibited by PCDH19-CE patients^18^. One study revealed the interaction of PCDH19 with N-cadherin, which regulates N-cadherin and β-catenin signalling, which is critical for neuronal and synaptic development^3^. Along with this, impairment in other basic mechanisms including excitatory synaptic contacts and neuronal activity^12,14^, and GABAergic transmission and inhibition^24,25^ were found as a result of mutations in *PCDH19*. In addition, there are several cellular PCDH19-CE models^7,26–28^. Studies, using human neurons reported abnormal cell sorting, cell migration, and asynchronized differentiation in neurons heterozygous for a mutation in *PCDH19* compared to WT cells^26,27^. These models also align with the cellular interference hypothesis proposed by Dibbens et al. 2008^5^.

In this study, we used CRISPR/Cas9 to generate a nonsense mutation knock-in mouse model of PCDH19-CE based on a similar mutation in a male patient with mosaic expression. We generated female (*Pcdh19^WT/mut^* and *Pcdh19^mut/mut^*) and male (*Pcdh19^mut^*) mice mutant for *Pcdh19*. RNA-sequencing of mouse brains from all the genotypes revealed genotype-specific gene expression patterns and suggests that there are neurodevelopmental pathways which are affected by the PCDH19 mutation. Furthermore, the expression of a specific set of genes was impaired only in the brains of heterozygous mutant females compared to WT and homozygous mutant females. Using neuronally differentiated human ESCs engineered to possess the same genetic mutation revealed that most of the genes that showed significant changes in expression in the heterozygous female brains in mice, showed a similar change in the human ESC differentiation system. Using the human ESCs neuronal differentiation assay in culture, we observed morphological differences between the different genotypes: Neurite length was significantly increased in PCDH19 mutant-heterozygous derived neurons and even further in neurons differentiated from PCDH19 mutant-homozygous ESCs. These findings suggest that some phenotypes may correlate with PCDH19 levels, while others are unique to the heterozygous genotype. Characterising and understanding these different phenotypes are important for deciphering the molecular basis for PCDH19-CE and the development of future therapy.

## Methods

### Mice and housing conditions

All in vivo experiments were approved by the Hebrew University Institutional Animal Care and Use Committee [#MD-24989-01] and were performed in accordance with the guidelines of the Hebrew University of Jerusalem, Institutional Animal Care and Use Committee (IACUC). All experiments were performed either on animals heterozygous or homozygous for *PCDH19* or on wildtype littermates. Animals were housed according to standard guidelines with free access to food and water in a 12hr light/dark cycle.

### Generation of mutant mice and mutant human Embryonic Stem cells

*Pcdh19*^WT/mut^ mice, *Pcdh19*^mut/mut^ and *Pcdh19*^mut^ (C57BL/6 background) were generated using CRISPR/Cas9 technology for generating the mutation c.T1548A, p.Y516*. Mutant mice were generated by replacing Tyrosine 516 with a termination stop codon in fertilized mouse oocytes, using CRISPR/Cas9 editing. Guide RNA (gRNA) (IDT) targeting the area of the desired mutation (c.T1548) was used. Recombinant CAS-9 protein was mixed with tracrRNA/gRNA to form a ribonucleoprotein (RNP) complex. This complex, combined with the donor DNA (50 µM), was electroporated into oocytes using the Neon transfection system (Invitrogen), according to the manufacturer’s guidelines. The modified blastocysts were then implanted into pseudo-pregnant female mice to generate F1 heterozygous mice. The mutation was verified by genotyping the progeny.

In stem cells, mutation, c.C1548A, p.Y516* was generated using CRISPR/Cas9 technology. The mutation was introduced using guide RNA (gRNA) (IDT) targeting the area of the desired mutation (c.C1548), which was pre-annealed to ATTO550-conjugated tracrRNA (IDT). Recombinant CAS9 protein was mixed with the tracrRNA/gRNA to form a ribonucleoprotein (RNP) complex. This complex, combined with the donor DNA (50 µM), was electroporated into 2×10^6^ WiBR3 human embryonic stem cells (hESCs) using the Neon transfection system (Invitrogen), according to the manufacturer’s guidelines. After electroporation, cells were plated with Nutristem media (Sartorius, cat. #05-100-1A), supplemented with the ROCK inhibitor Y-27632 (Enco, cat. #10005583) and an HDR enhancer (homologous recombination enhancer, IDT, cat. #10007910). Media was replaced daily. 48 hours after electroporation, cells were trypsinized and sorted by FACS for ATTO550-positive cells. Sorted cells were seeded on mouse embryonic fibroblasts (MEF)-coated plates with ROCK inhibitor. 4-5 days later, individual clones were picked and replated in 24-well plate. After expansion of the clones, each clone was split to a Matrigel-coated plate and grown further for DNA, RNA and protein extraction. Presence of the C1548A mutation was verified by Sanger sequencing. PCDH19 expression levels were measured by RT-Q-PCR and Western blot.

### Preparation of mouse embryonic fibroblasts

MEFs were cultured in DMEM media containing 15% FBS, 1% P/S, 1% L-Glutamine, 1% NEAA (non-essential amino acids) and 50 µM β-mercaptoethanol. MEFs were trypsinized and irradiated at 30 Gy and subsequently stored in liquid nitrogen. Plates were coated with autoclaved 0.2% gelatin (Sigma, cat. #G2500) in molecular grade water for 10 minutes. Irradiated MEFs were thawed and plated on the coated plates, at least 24 hours prior to plating hESCs.

### Human Embryonic Stem cells (hESCs)

WiBR3 hESCs (from female source) were grown on MEF feeder layer in DMEM/F12 (Sartorius, cat. #01-170-1A) medium supplemented with 15% knockout serum replacement (KOSR) (Gibco, cat. #10828-028), 1% NEAA, 1% P/S, 1% sodium pyruvate, 50 µM β-mercaptoethanol and 8 ng/ml Fibroblast Growth Factor-2 (FGF-2) (Peprotech cat. #100-18B). Cell cultures were supplemented with ROCK inhibitor (Y27632 final concentration 10 µM) (Enco, cat. #10005583) during passaging to support seeding of single cells. Media was changed daily. For passaging, cells were trypsinized with trypsin C when cultured on MEFs (Biological Industries, cat. #03-053-1B). For extraction of RNA, genomic DNA, proteins, infection and differentiation, cells were seeded on Matrigel (1:120, Corning, cat. #356237), without the MEF feeders, in mTser1 media (Stem Cell Technologies, cat. #85850) and ROCK inhibitor (Y27632 final concentration 10 µM). Media was changed daily and cells were passaged upon reaching 70%-80% confluency. When cells were cultured on Matrigel, Accutase (Sigma, cat. #SCR005) was used for passaging. All cells were maintained at 37°C, 5% CO_2_. To freeze the cells, ACF, Protein-free NutriFreez D10 Cryopreservation Medium (Biological Industries, cat. # 05-713-1B) was used.

### Virus preparation

DNA mix containing 1 µg lentiviral vector, 0.5 ug pMD2.G (Addgene, cat. #12259) and 0.5 µg psPAX2 (Addgene, cat. #12260) was prepared. Mix containing 100 µl OptiMEM (Gibco, cat. #51985034) and 6 µl of transfection reagent FuGENE-HD (Promega, cat. #609920) was added to the DNA mix and incubated for 20 minutes at room temperature (RT). After incubation, the transfection mix was added to cell suspension containing 1.25×10^6^ HEK293T cells and seeded in a 6-well plate. The day after transfection, medium was replaced with 2.5 ml fresh medium. The day after (48 hours post transfection), medium containing viruses was collected, filtered through a 0.45 mm filter to remove cell debris and aliquoted. The virus was then concentrated using PEG-8000 at a ratio of 1:4 (Sigma, cat. #81268) and incubated at 4°C for 30 minutes to overnight. The mixture was centrifuged at 1500g for 45 minutes at 4°C. The virus was then resuspended in mTser media and either used immediately or stored at -80°C.

### Infection and selection

400,000 hESCs/well were plated on Matrigel-coated 6 well plate. The cells were infected with a 1:4 virus dilution, in a volume of 1.5 ml. Polybrene was added to a final concentration of 8 µg/ml. The next day, medium was changed to 2 ml fresh medium. The next day (48 hours post infection), selection with Puromycin at 2 µg/ml was initiated. Selection media was replaced every two days until the end of selection, as determined by complete cell death of the control uninfected cells.

### Differentiation of hESCs into cortical neurons

Neuronal differentiation protocol was adopted and modified from Zhang et al^29^. 1×10^6^ ESCs were plated on Matrigel-coated 6 well plate with mTser and ROCK inhibitor. The next day, media was replaced by Neuronal Induction Media (DMEM/F12 with HEPES, 1% penicillin/streptomycin, 1% L-glutamine, 1% 100X N2 supplement (Thermofisher, cat. #17502048) and 1% 100X NEAA) and freshly added 2µg/ml doxycycline. The media was replaced every two days until we observed a significant morphology change. Simultaneously, PDL (Sigma, cat. #1024)/Laminin (GIBCO, cat. #23017-015)-coated plates were prepared. After the cells resembled neural progenitors, they were detached using Accutase and 500,000 cells/well were seeded on 12 well plates coated with PDL/laminin in Cortical Induction media (DMEM/F12 with HEPES, Neurobasal Media (Rhenium, cat. #21103-049), 2% of 50x B27 supplement (GIBCO, cat. # 17504044), 1% penicillin/streptomycin, 0.1% Brain Derived Neurotrophic Factor (Abcam, cat. # AB9794), 0.1% Glial Derived Neurotrophic Factor (Peptrotech, cat. #450-10), 0.1% NT-3 (Abcam, cat. # AB9792), 0.1% laminin (Gibco, cat. #23-017-015), 2µM AraC (Sigma, cat. # C6645) and freshly added 2µg/ml doxycycline. Half of the cortical neural media was replaced every 4 days and cells were kept in culture for 10 days. Cells were then harvested for RNA, and protein, and fixed for imaging.

### RNA Sequencing-Sample Preparation

Three mice from each genotype, *Pcdh19*^WT/WT^ female, *Pcdh19*^WT/mut^ female, *Pcdh19*^mut/mut^ female, *Pcdh19*^WT^ male and *Pcdh19*^mut^ male, were sacrificed in accordance with the guidelines of the Hebrew University of Jerusalem Institutional Animal Care and Use Committee (IACUC). RNA was isolated using TRI-Reagent (Sigma, cat. #T9424) from whole brain tissue for each sample and was quantified using Nanodrop. 2µg of RNA was sent for sequencing. Sequencing was performed using Novaseq.

### RNAseq Analysis

Raw reads were aligned to the mouse transcriptome and genome version GRCh39 with annotations from Ensembl release 106 using TopHat v2.1.1^30^. Counts per gene quantification was done using htseq-count v2.01^31^. Genes with a sum of counts below 10 over all samples were filtered out. Normalization and differential expression analysis were performed with the DESeq2 package v 1.36.0^32^. Differentially expressed genes (DEGs) were defined by applying a significance threshold of *\*P* <0.05 and Log2FC < -0.585 or > 0.585.

### Enrichment Analysis

Gene Ontology enrichment analysis, molecular function, and biological process was performed using EnrichR (https://maayanlab.cloud/Enrichr/) with p-val<0.05, Log2FC<-0.585 or >0.585.

### Reverse Transcription PCR (RT-PCR)

RNA was extracted and converted to cDNA using iScript cDNA Synthesis Kit (Bio-Rad, cat. #1708891). PCR was performed using the 2X Taq HS Mix Red PCR MasterMix (PCR Biosystems, cat. #PB10.23.02) according to the manufacturer’s guidelines. Annealing was performed at 60 °C and 35-40 cycles used to amplify targets. PCR products were visualized on a 2% agarose gel with ethidium bromide (EtBr) and visualized on Gel Doc XR+ imaging system using ImageLab software (Bio-Rad).

### Quantitative Reverse Transcription-PCR (RT-Q-PCR)

RT-Q-PCR was performed on the cDNA using iTaq Universal SYBR Green Supermix (Bio-Rad), and the CFX96/384 (Bio-Rad) real-time PCR machine. Normalization was performed using primers for the house keeping genes *HPRT, NDU or GAPDH* (unless mentioned otherwise). Primers were calibrated using serial dilutions of cDNA to validate their efficiency. ΔΔCt method was used to calculate the gene expression changes. Expression levels show relative-fold change compared to control samples, which was set to 1 for general understanding. RT-Q-PCR was repeated in triplicates.

### Protein extraction and quantification

Protein was extracted from cells and tissue using RIPA Lysis Buffer. RIPA constituted of 50mM Tris-HCl pH8, 150mM NaCl, 1% Triton-X-100, 0.1% SDS, 0.5% Sodium Deoxycholate and one protease inhibitor cocktail tablet (Roche, cat. #11836153001) for every 10 ml RIPA. The solution was stored at 4°C or aliquoted and stored at -20°C. Cold RIPA was added to the cells or tissue and were either scraped (cells) or homogenized using magnetic beads (tissue) until the solution is homogenous. Then the solution was incubated on ice for 30 minutes with quick vortex every 10 minutes before it was centrifuged at 14000 rpm for 15 minutes at 4°C. The clear lysate was transferred to new tubes for protein quantification. Protein quantification was performed using DC Protein Assay (Bio-Rad, cat. #500-0111) according to the manufacturer’s guidelines. Absorbance was calculated using the Bio-Rad microplate reader at 750 nm. Bovine Serum Albumin was used to obtain a protein standard curve.

### Western Blotting

SDS-polyacrylamide gel electrophoresis (SDS-PAGE) was used to separate the proteins. 30-50 µg of protein was loaded on each well, unless otherwise stated, on 10% polyacrylamide gel using the Mini-PROTEAN Gel Electrophoresis apparatus at 120 mV for approximately 2 hours. Gels were then transferred to PVDF membrane (Merck Millipore) in a 20% methanol transfer buffer using a transblot machine (Bio-Rad) with the standard program, followed by the high molecular weight program because the PCDH19 protein has a molecular weight of 135 kD. After the transfer, blocking was performed in 5% skim milk in TBST, followed by washes and incubation with either anti-PCDH19 (Bethyl Laboratories, cat. # A304-468A, 1:1000), anti-GAPDH (Sigma, cat. #015M4824V, 1:5000), anti-Tubulin (Abcam, cat. #Ab6160, 1:1000), or anti- β-catenin (BD Biosciences, cat. #610154, 1:1000) for 1 hour at RT or at 4°C overnight. After primary antibody incubation, membranes were washed with TBST and incubated with respective secondary antibody at a dilution of 1:5000 in 5% skim milk in TBST for 40-50 minutes at RT. After washing the secondary antibody, detection was performed using ECL on a Gel Doc XR+ imaging system and ImageLab software (Bio-Rad).

### Immunofluorescence

Cells were seeded either on coverslips in a 12-well plate or u-slide 8 well IbiTreat (Ibidi, cat. #80826) chambers at appropriate confluency. Cells were washed with PBS and fixed using 4% paraformaldehyde (PFA) at the desired time point. The cells were incubated with PFA for 10 minutes followed by three washes with PBS. For permeabilization, ice cold 70% ethanol was added and the cells were incubated at -20°C overnight or longer. The next day, blocking was performed using 5% BSA/ PBST (PBS supplemented with 0.1% Tween20) and incubated for 1 hour at RT. After blocking, desired volume of primary antibody, anti-tubulin β3 (Biolegend, cat. #801201, 1:1000), anti-PCDH19 (Bethyl Laboratories, cat. # A304-468A, 1:1000), anti-GFAP (Abcam, cat. #4674), diluted in blocking solution, was added and samples were incubated either for 2 hours at RT or overnight at 4°C. Then cells were washed with PBS and suitable secondary antibody, diluted in blocking solution, was added and incubated for 1 hour at RT. Cells were then washed again and mounted using DAPI mounting media (Vectashield, cat. #H-1200). After drying, cover slips were sealed using transparent nail polish. Slides were kept in the dark 4°C until imaging. Confocal microscopy images were taken using NIKON spinning disk confocal microscopy and processed via NIS-elements software.

### Confocal microscopy and Image Analysis

Images were taken using NIKON TS100 microscope at 10X, 20X and 40X magnification and viewed using NIKON digital sight device and NIS-elements software and analyzed using Imaris Software and QuPath.

### Neurite length assay

Neurite length was calculated using Imaris Software. The total neurite length was calculated in the image and were averaged to the total number of cells, between 100-500 cells for each genotype.

## Results

### Generation of *Pcdh19*-mutant mouse model

Most of the mutations that cause PCDH19-CE are located in exon 1 of the *PCDH19* gene. In order to investigate the molecular mechanism of PCDH19-CE, we used CRISPR/CAS9 to knock-in a nonsense *Pcdh19* mutation into mice, identical to the mutation in one of the patients (c.T1548A, p.Y516*). This mutation, Thymine (T) to Adenine (A) at nucleotide 1548, results in a premature termination codon (PTC) (**Fig. 1A**). Sanger sequencing of genomic DNA isolated from female mice was performed to confirm the genetic mutation. We observed that along with a female mouse heterozygous for the mutation (*Pcdh19*^WT/mut^), we obtained a female mouse which was homozygous for the mutation (*Pcdh19*^mut/mut^) (**Fig. 1B**). Furthermore, the presence of the mutation was confirmed by genotyping the progeny using wildtype (WT) and mutation (mut) specific PCR primers (**Supplementary Fig. 1**). Male mice hemizygous for the mutation (*Pcdh19*^mut^) were also obtained and validated (**Fig. 1B**). Using Western blot analysis, we observed that PCDH19 protein expression level was reduced by approximately half in heterozygous *Pcdh19*^WT/mut^ mice compared to *Pcdh19*^WT/WT^ mice and undetectable in homozygous *Pcdh19*^mut/mut^ and hemizygous *Pcdh19*^mut^ mice (**Fig. 1C, D).**

**Figure 1.**
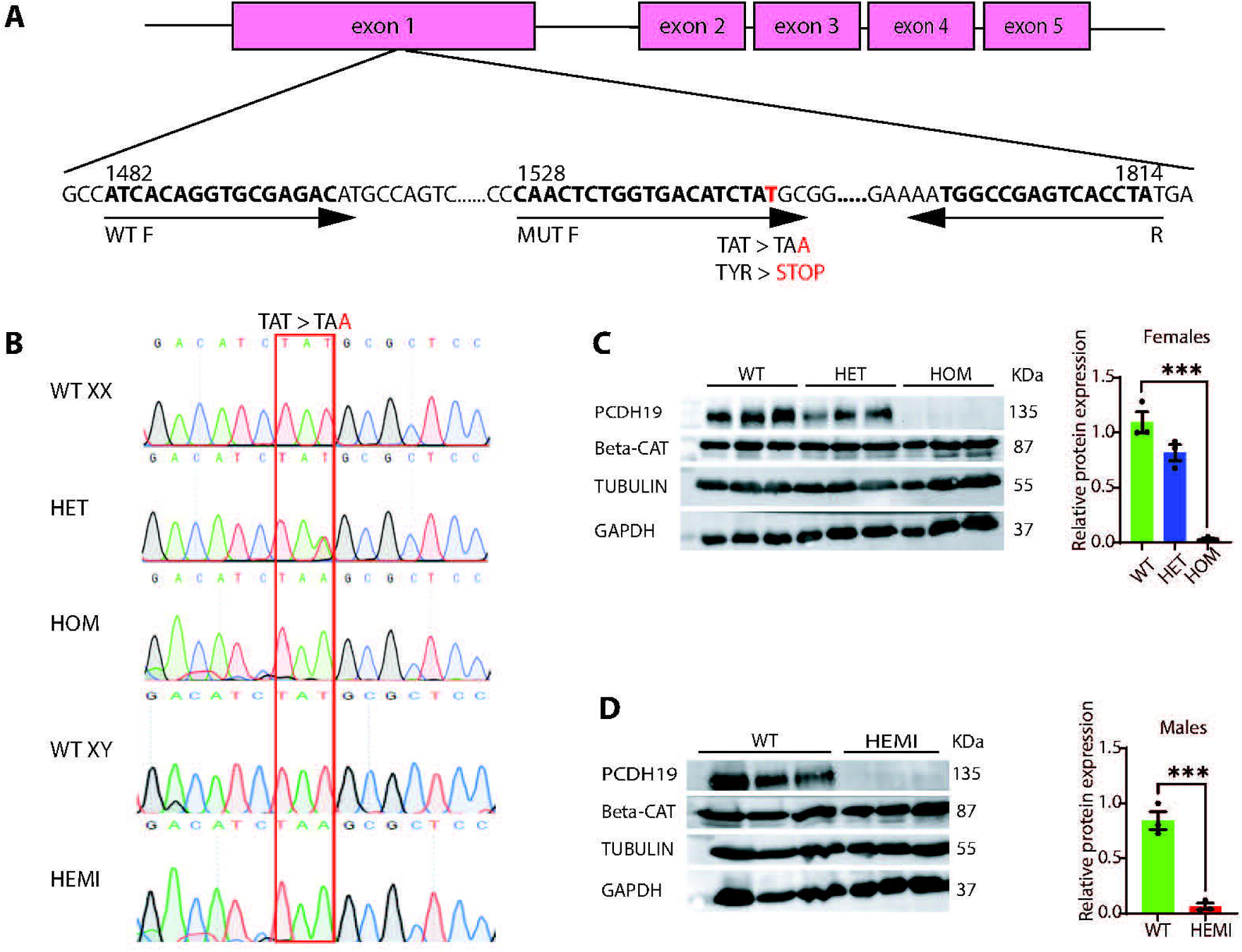
Generation of Pcdh19-mutant mice using CRISPR-Cas9. (**A**) Scheme for generating mutant mice model for PCDH19-CE. The mutation p.Y516* was introduced into Pcdh19 exon 1 using CRISPR. The mutation leads to premature termination codon causing the transcript to undergo nonsense mediated decay. (**B**) Sanger sequencing of Pcdh19 ^WT/WT^ (WT), Pcdh19 ^WT/mut^ (HET) and Pcdh19^mut/mut^ (HOM) female mice, and Pcdh19 ^WT^ (WT) and Pcdh19^mut^ (HEMI) male mice. (**C**) Western blot analysis of protein lysates (left) and quantification of PCDH19 protein (right) from brains of Pcdh19 ^WT/WT^ (WT), Pcdh19 ^WT/mut^ (HET) and Pcdh19^mut/mut^ (HOM) female mice. Unpaired T-test: ***p = 0.008. (D) Western blot analysis of protein lysates (left) and quantification of PCDH19 protein (right) from brains of Pcdh19 ^WT^ (WT) and Pcdh19 ^mut^ (HEMI) male mice. n = 3 mice per group. Unpaired T-test: ***p<0.005.

### Gene Expression Profile of *Pcdh19*-mutant Mice

To investigate the molecular differences between *Pcdh19* mutant mice and WT mice, we isolated RNA from brains of mice aged 8-10 weeks and performed RNA-sequencing (n=3, per genotype). Principal component analysis (PCA) of the RNA-sequencing data identified that gene expression patterns of *Pcdh19*^WT/WT^ and *Pcdh19*^mut/mut^ female mice clustered together, while gene expression pattern of *Pcdh19*^WT/mut^ female mice clustered separately (**Fig. 2A**). Gene expression analysis of all samples identified 186 significantly differentially expressed genes (DEGs) (*\*P* <0.05; Log2FC < -0.585 or >0.585) between *Pcdh19*^WT/WT^ and *Pcdh19*^WT/mut^ mice (**Fig. 2B, C**) and 125 DEGs between *Pcdh19*^WT/mut^ and *Pcdh19*^mut/mut^ mice (**Fig. 2B, D**). MA plot, scatter plot and volcano plots show the distribution of DEGs in both the *Pcdh19*^WT/WT^ vs *Pcdh19*^WT/mut^ comparison and the *Pcdh19*^WT/mut^ vs *Pcdh19*^mut/mut^ comparison. While these comparisons revealed a large number of DEGs, each red dot corresponds to a gene which is differentially expressed, a comparison of gene expression of *Pcdh19*^WT/WT^ vs *Pcdh19*^mut/mut^ mice did not yield many DEGs (**Supplementary Fig. 2A-I**). Overlap of DEGs from *Pcdh19*^WT/WT^ vs. *Pcdh19*^WT/mut^ comparison and *Pcdh19*^WT/mut^ vs. *Pcdh19*^mut/mut^ comparison identified a set of 33 shared genes (**Fig. 2B**). Out of these 33 genes, 30 were downregulated specifically in *Pcdh19*^WT/mut^ mice, 3 were upregulated specifically in *Pcdh19*^WT/mut^ mice (marked in red) and 2 genes (*Eno1b* and *Pcdh19*) were downregulated in both *Pcdh19*^WT/mut^ and *Pcdh19*^mut/mut^ mice. Expression of *Eno1B*, was non-uniform and thus it was excluded from further analyses. The Venn diagram shows an overlap of 33 genes between the two comparisons (**Fig. 2B**).

**Figure 2.**
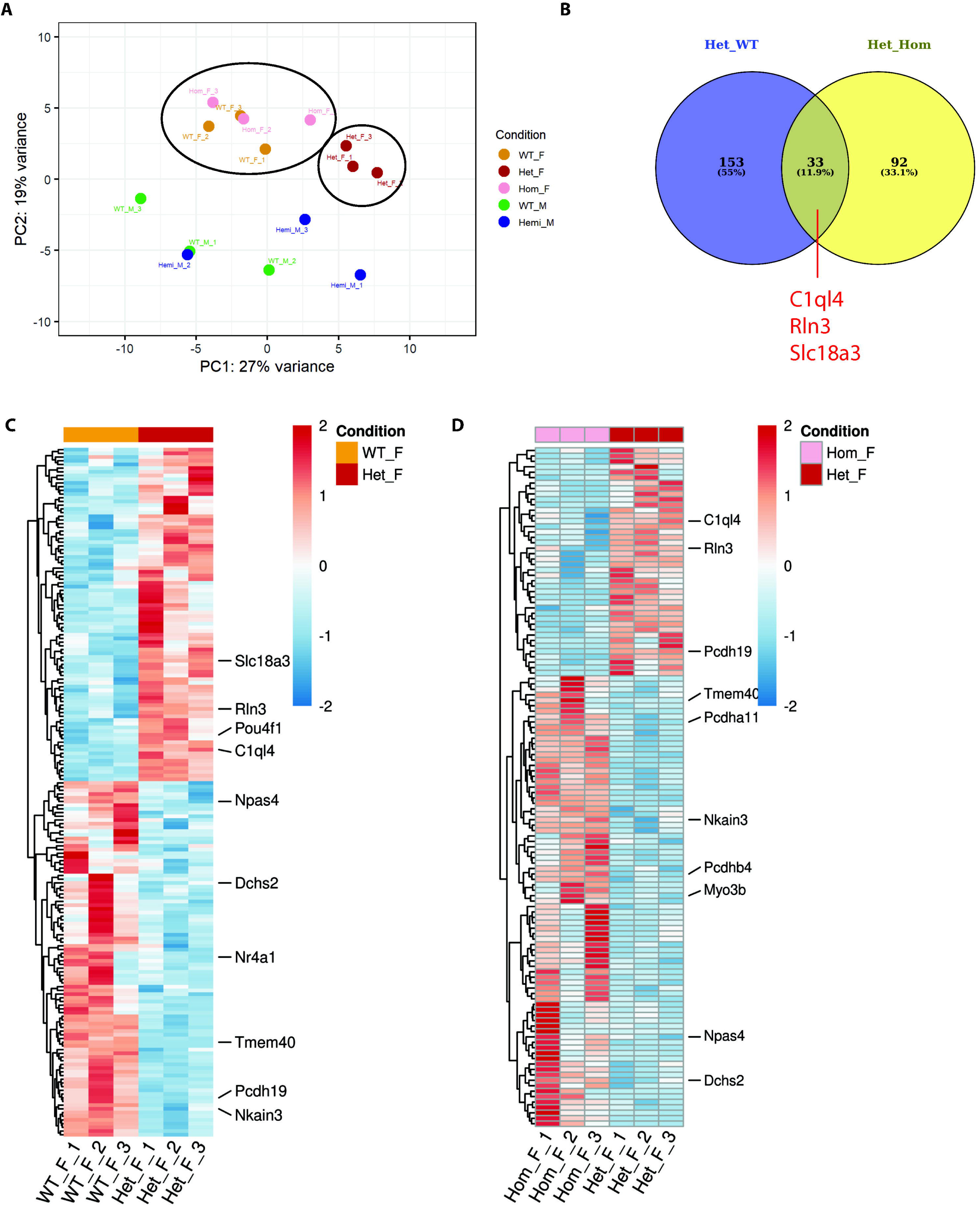
RNA sequencing of brains of Pcdh19-mutant mice reveal differentially expressed genes (DEGs) (**A**) Principal Component Analysis of RNA sequencing data from Pcdh19 ^WT/WT^ female mice (WT_F), Pcdh19 ^WT/mut^ female mice (Het_F), Pcdh19^mut/mut^ female mice (Hom_F), Pcdh19 ^WT^ male mice (WT_M) and Pcdh19 ^mut^ male mice (Hemi_M) (n=3 mice per genotype). The figure features a three-dimensional scatter plot of the principal components (PCs) of the data. Each point represents an RNA-Seq sample. Samples with similar gene expression profiles are clustered together. Sample groups are indicated by different colors. (**B**) Venn Diagram representing the overlap of DEGs (33) between Pcdh19 ^WT/mut^ vs. Pcdh19 ^WT/WT^ (Het_WT, 186) and Pcdh19 ^WT/mut^ vs. Pcdh19 ^mut/mut^ (Het_Hom, 125) comparisons based on the criteria, p-val <0.05 and Log2(FC)<-0.585 or >0.585. 30 of the overlapping genes were downregulated in Pcdh19 ^WT/mut^ mice compared to Pcdh19 ^WT/WT^ and Pcdh19 ^mut/mut^ mice. Only three (marked in red) were upregulated. (**C**) Heat map representing DEGs between Pcdh19 ^WT/WT^ female (WT_F) and Pcdh19 ^WT/mut^ female (Het_F) mice. Each row represents a gene, each column represents a sample and each cell displays normalized gene expression values. The Log2(FC) is represented by the color bar on the right. Genes of interest are labeled. (D) Heat map representing DEGs between Pcdh19 ^mut/mut^ female (Hom_F) and Pcdh19 ^WT/mut^ female (Het_F) mice. Each row represents a gene, each column represents a sample and each cell displays normalized gene expression values. The Log2(FC) is represented by the color bar on the right. Genes of interest are labeled.

### Identification of DEGs and Pathway Enrichment in *Pcdh19*-Mutant Mice

Focusing on the list of 186 DEGS from the *Pcdh19*^WT/WT^ vs. *Pcdh19*^WT/mut^ comparison and 125 DEGs from *Pcdh19*^WT/mut^ vs. *Pcdh19*^mut/mut^ comparison, we performed pathway enrichment analysis. The analysis was performed separately for genes that were either upregulated in *Pcdh19*^WT/mut^ mice or downregulated in *Pcdh19*^WT/mut^ mice for each comparison. We found enrichment of genes in pathways involved in activation of Hox genes during differentiation, synaptic transmission, transmembrane export, inhibitory postsynaptic potential and others in *Pcdh19*^WT/mut^ upregulated genes (**Fig. 3A)**. In *Pcdh19*^WT/mut^ downregulated genes we detected enrichment of genes in NTRK signalling, AKT phosphorylation, binding of TCF/LEF promoters and NPAS4 mediated regulation of genes and voltage-gated potassium channel activity pathways (**Fig. 3B).** Similar pathways were enriched in a comparison of *Pcdh19*^WT/mut^ vs. *Pcdh19*^mut/mut^ DEGs. DEGs upregulated in *Pcdh19*^WT/mut^ mice, in a comparison of *Pcdh19*^WT/mut^ vs. *Pcdh19*^mut/mut^ DEGs, were enriched for pathways such as phenylalanine and tyrosine metabolism, zinc transporters and influx, estrogen biosynthesis and relaxin receptors. DEGs downregulated in *Pcdh19*^WT/mut^ mice were enriched for pathways that included respiratory metabolism, voltage-gated potassium channels, RUNX and AKT signalling (**Fig. 3C, D).**

**Figure 3.**
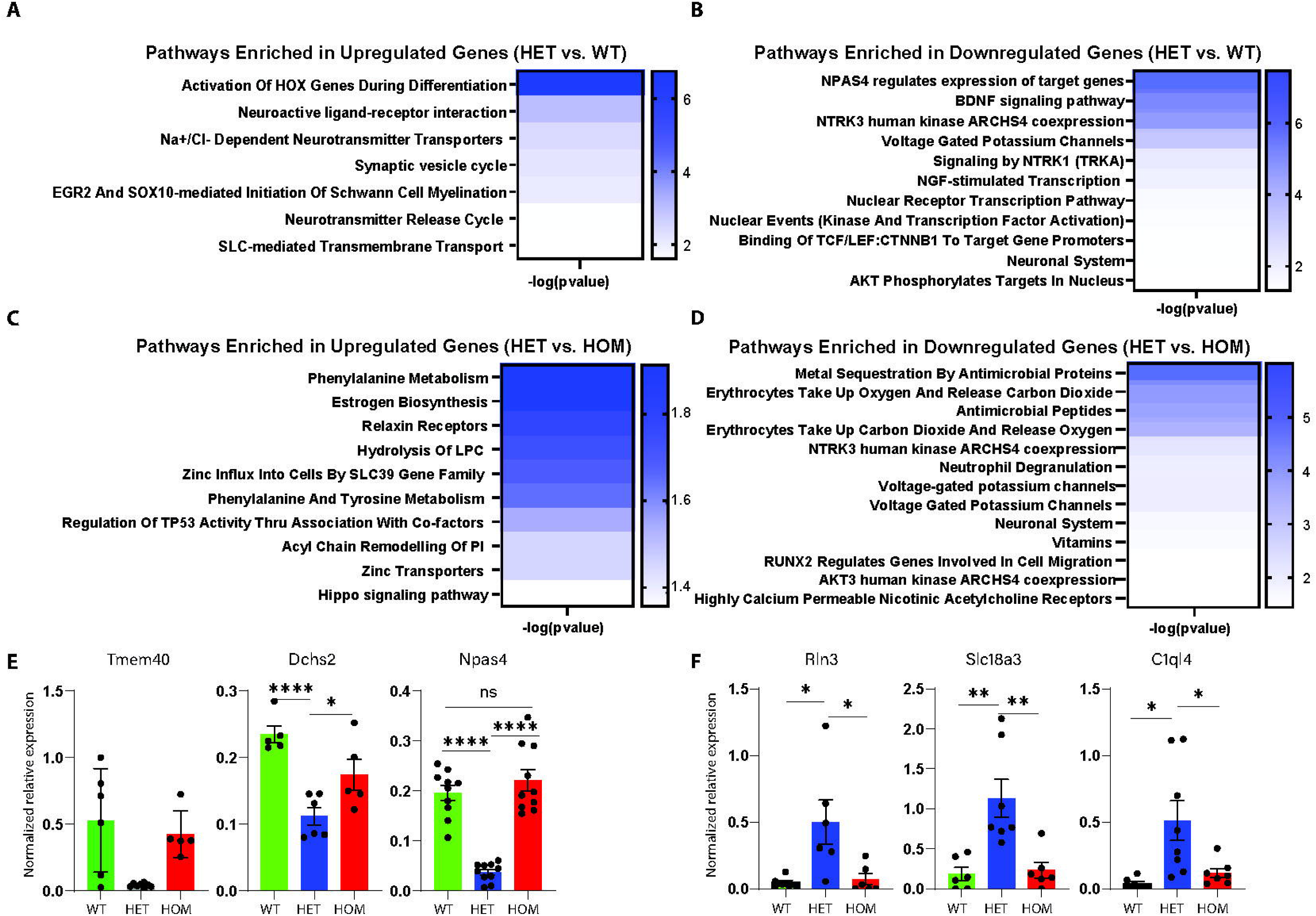
Pathway and functional enrichment analysis of DEGs in Pcdh19-mutant mice brain samples. (**A**) Enriched pathways based on DEGs upregulated in Pcdh19 ^WT/mut^ in Pcdh19 ^WT/WT^ (WT) vs. Pcdh19 ^WT/mut^ (HET) comparison. (**B**) Enriched pathways based on DEGs downregulated in Pcdh19 ^WT/mut^ in Pcdh19 ^WT/WT^ (WT) vs. Pcdh19 ^WT/mut^ (HET) comparison. (**C**) Enriched pathways based on DEGs upregulated in Pcdh19 ^WT/mut^ in Pcdh19 ^WT/mut^ (HET) vs. Pcdh19 ^mut/mut^ (HOM) comparison. (D) Enriched pathways based on DEGs downregulated in Pcdh19 ^WT/mut^ in Pcdh19 ^WT/mut^ (HET) vs. Pcdh19 ^mut/mut^ (HOM) comparison. (E) Validation of genes downregulated in Pcdh19 ^WT/mut^ (HET) female mice by RT-Q-PCR on mice brain samples from Pcdh19 ^WT/WT^ (WT), Pcdh19 ^WT/mut^ (HET) and Pcdh19 ^mut/mut^ (HOM) female mice (n = 5-7 mice per group). Unpaired T-test: *p<0.01, **p<0.005, ****p<0.0001. Additional genes are presented in Supplementary Fig. 3. (**F**) Validation of genes upregulated in *Pcdh19* ^WT/mut^ using RT-Q-PCR on mice brain samples *Pcdh19* ^WT/WT^ (WT), *Pcdh19* ^WT/mut^ (HET) and *Pcdh19* ^mut/mut^ (HOM) female mice (n = 5-7 mice per group). Unpaired T-test: *p<0.01, **p<0.005, ****p<0.0001. Additional genes are presented in Supplementary Fig. 3.

Focusing on the set of 33 overlapping genes that are differentially expressed in both the *Pcdh19*^WT/WT^ vs. *Pcdh19*^WT/mut^ comparison and the *Pcdh19*^WT/mut^ vs. *Pcdh19*^mut/mut^ comparison, we validated the expression of several genes by RT-Q-PCR. In agreement with the RNA-sequencing data, we observed lower mRNA expression of *Tmem40*, *Dchs2*, and *Npas4* in *Pcdh19*^WT/mut^ mice brains compared to *Pcdh19*^WT/WT^ and *Pcdh19*^mut/mut^ mice brains, and higher mRNA expression of *Rln3*, *C1ql4* and *Slc18a3* in *Pcdh19*^WT/mut^ mice brains compared to *Pcdh19*^WT/WT^ and *Pcdh19*^mut/mut^ mice brains (**Fig. 3E, F**). Additional DEGs were validated, including immediate early genes (*Erg2, Arc* and *Nr4a1)* and *Hox* genes, many of which showed down- or up-regulation in brains of both heterozygous and homozygous mice (**Supplementary Fig. 3A, B**). Pathway enrichment analysis and GO analysis on 30 of the downregulated genes in *Pcdh19*^WT/mut^ out of the 35 overlapping DEGs in *Pcdh19*^WT/WT^ vs. *Pcdh19*^WT/mut^ comparison and the *Pcdh19*^WT/mut^ vs. *Pcdh19*^mut/mut^ comparison, identified pathways involved in NTRK signalling, neuronal system, NMDA mediated transmission, long-term potentiation, synaptic transmission, and inhibitory post-synaptic potential (**Supplementary Fig. 3C**).

### Generation of human *PCDH19*-mutant ESC cellular model

We next wanted to establish a cellular human neuronal model system (**Fig. 4A**). For this purpose, we generated the mutation (c.C1548A, p.Y516*) in a hESC line (WibR3, female) using CRISPR/CAS9 (**Fig. 4**). Individual cell clones were isolated and Sanger sequencing confirmed clones both homozygous and heterozygous for the mutation (**Fig. 4B**). We transduced these cells with the NGN2/ASCL1 tet-inducible system^33^. Following tet induction and neural induction of WT, homozygous and heterozygous *PCDH19*-mutant ESCs, we monitored differentiation by mRNA expression levels of stem cell markers (*SALL* and *OCT4*) and neuronal markers (*TUBB3* and *MAP2*) at day 0 and 12 of the differentiation protocol. We observed an increase in mRNA expression of neuronal markers (*TUBB3* and *MAP2*) and a concomitant decrease in mRNA expression of stem cell markers (*OCT4* and *SALL*) on day 0 and 12 (**Fig. 4C**). We also determined the *PCDH19* mRNA and protein levels in the differentiated neurons. As expected, *PCDH19* mRNA levels reduced to approximately 60 and 20 percent in heterozygous and homozygous PCDH19-mutant clones, respectively (**Fig. 4D-E)**.

**Figure 4.**
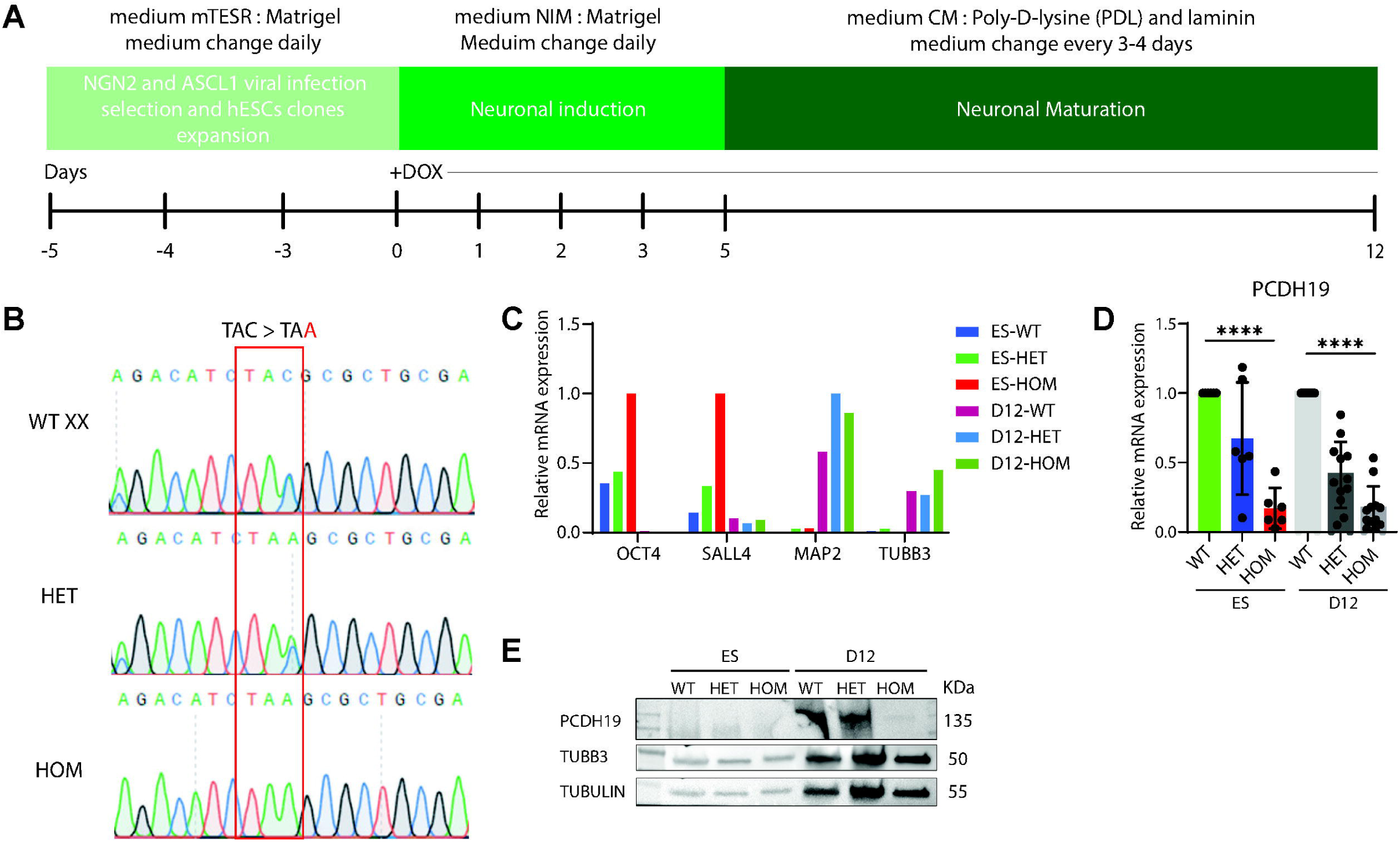
In vitro human neuronal model for PCDH19-CE. (**A**) Schematic of the generation of induced neurons from human ES cells. (**B**) Verification of genotypes of wildtype (WT), heterozygous (HET) and homozygous (HOM) PCDH19-mutant ES cell clones generated by CRISPR-CAS9. (**C**) RT-Q-PCR of RNA from human ES cells at day 0 and 12 of neuronal induction for ES markers (SALL and OCT4) and neuronal markers (TUBB3 and MAP2) (n = 3 biological repeats). Unpaired T-test: ****p<0.0001. (D) Gene expression levels of PCDH19 from wildtype (WT), heterozygous (HET) and homozygous (HOM) PCDH19-mutant clones at day 0 and day 12 of neuronal induction using RT-Q-PCR (n > 5 biological repeats). Unpaired T-test: ****p<0.0001. (E) Western blot analysis of protein lysates of PCDH19 protein from wildtype (WT), heterozygous (HET) and homozygous (HOM) human PCDH19-mutant ES clones at day 0 and day 12 of neuronal induction.

### Characterization of the cellular phenotype of human *PCDH19-*mutant induced neurons

Having established a human *PCDH19*-mutant cellular model, we next wanted to determine if there is a phenotypic difference between neurons induced from the *PCDH19*-mutant clones (homozygous and heterozygous) compared to WT. For this purpose, *PCDH19*-mutant and WT ESCs were differentiated and fixed at day 10. Cells were stained for neuronal marker TUBB3 and DAPI. We observed that heterozygous and homozygous *PCDH19*-mutant induced neurons had significantly longer neurite lengths than WT induced neurons, with homozygous *PCDH19*-mutant derived neurons exhibiting the longest neurites (**Fig. 5A, B**). Thus, there is a clear phenotype of longer neurite lengths in the human *PCDH19*-mutant derived neurons which inversely correlates with WT PCDH19 levels. This is in contrast to what we observed in mouse brain gene expression analysis, where the *Pcdh19*^mut/mut^ genotype showed similar patterns of gene expression as the WT mice (**Fig. 3E-F**).

**Figure 5.**
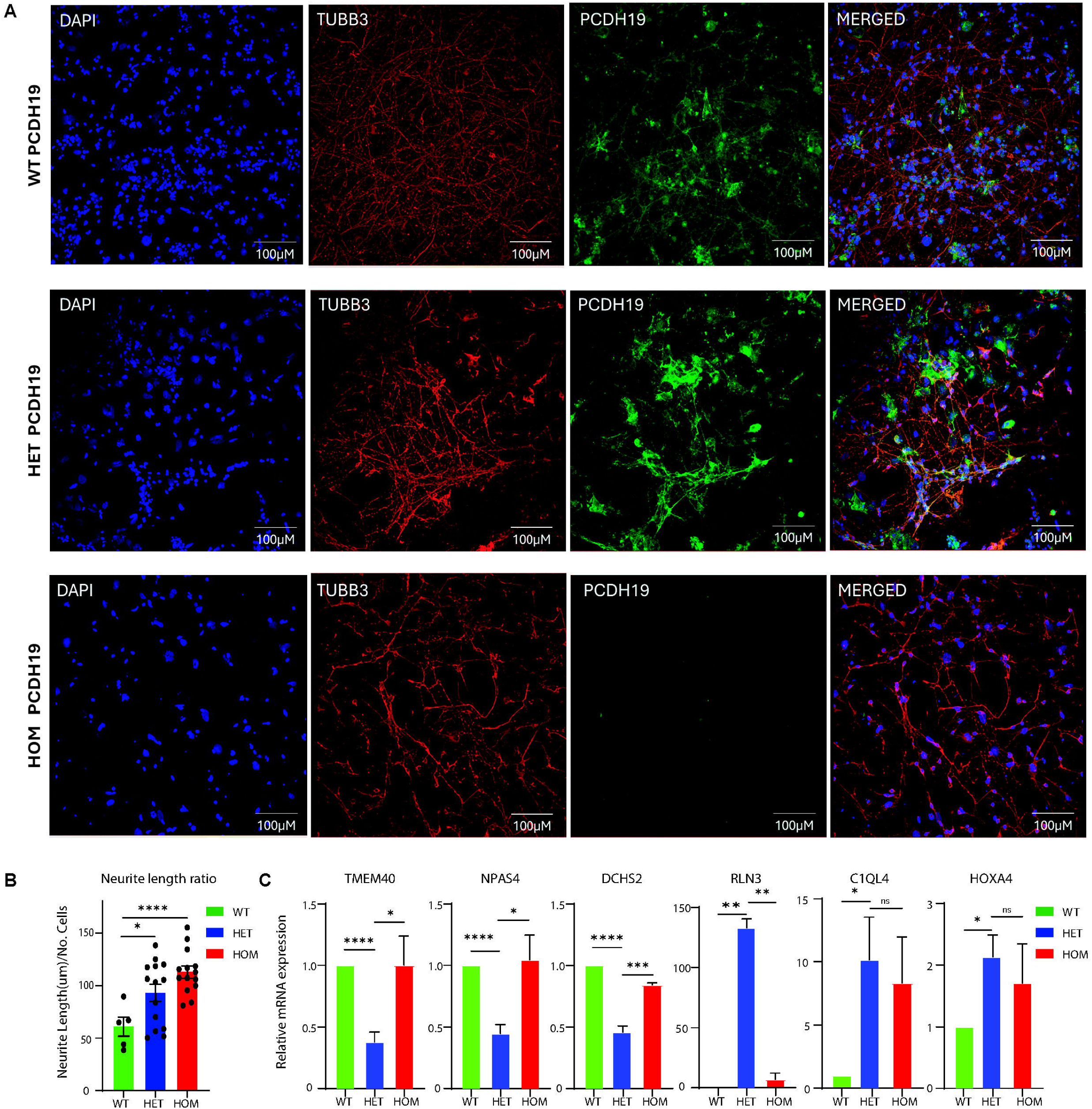
Characterization of phenotype of in vitro human neuronal model for PCDH19-CE. (**A**) Immunofluorescence of day 10 cortical neurons from wildtype (WT), heterozygous (HET) and homozygous (HOM) PCDH19-mutant clones stained with PCDH19 (green), Tubb3 (red), and DAPI (blue). (**B**) Neurite length for different genotypes was measured using Imaris (n= 3 biological replicates). Unpaired T-test: ****p<0.0001. (**C**) RT-Q-PCR of DEGs identified from Pcdh19-mutant mice brains on human wildtype (WT), heterozygous (HET) and homozygous (HOM) PCDH19-mutant induced neurons at day 12 (n= 4-9 per group, n=2 for RLN3). Unpaired T-test: *p<0.05, ***p<0.005, ****p<0.0001

We therefore wanted to examine the expression levels of the DEGs, that were identified in the RNA sequencing analysis of mice brain samples, in *PCDH19*-mutant and WT human ESCs differentiated clones. Similar to what we observed in mouse brains (**Fig. 3**), *TMEM40*, *NPAS4*, and *DCHS2* mRNA expression was lower in the differentiated heterozygous *PCDH19*-mutant clone compared to the differentiated WT and homozygous *PCDH19*-mutant clones, while mRNA expression of *C1QL4* and *RLN3* was higher in differentiated heterozygous *PCDH19*-mutant clones compared to differentiated WT and homozygous *PCDH19*-mutant clones. HOXA4 was upregulated in both differentiated heterozygous and homozygous PCDH19-mutant clones relative to WT (**Fig. 5C**). These observations are consistent with the *Pcdh19*-mutant and WT mouse brain RNA sequencing results. Expression of additional genes (*ARC, NR4A1, C-FOS, EGR2, NKAIN3SLC18A3*) were also validated in human neurons, however, these genes showed an opposite pattern to what was observed in the mouse model (**Supplementary Fig. 4**). Altogether, these results support the involvement of these genes in manifesting the PCDH19-CE phenotype.

## Discussion

Mutations in the *PCDH19* gene manifests a severe neurological phenotype in heterozygous females. Most patients do not respond to anti-epileptic medications, highlighting the need to understand the underlying molecular pathological mechanisms of PCDH19-CE in order to develop effective treatments for this unique disorder. In this work, we generated and characterized both a mouse model, as well as, a human cellular model harbouring a nonsense *PCDH19* mutation. RNA sequencing of brain samples from mice with different *Pcdh19* mutant genotypes identified DEGs and enriched pathways in *Pcdh19*^WT/mut^ heterozygous mice compared to *Pcdh19*^mut/mut^ homozygous and WT mice (**Figs 2, 3**). The transcriptional signatures from mouse brains, reveal for the first time that homozygous *PCDH19*-mutant neuronal-like cells exhibit molecular profiles more similar to WT cells than to heterozygous *PCDH19*-mutant cells, further supporting the cellular interference hypothesis^34^. Most of the DEGs identified in the mouse model were validated using human ESCs, either homozygous or heterozygous for the *PCDH19* mutation, following induction of neural differentiation (**Fig. 5C**).

Using human induced neuronal-like cells, we characterized a cellular phenotype specific to neurons carrying the *PCDH19* mutation. Both heterozygous and homozygous *PCDH19*-mutant neuronal-like cells exhibited significantly longer neurite lengths compared to WT neuronal-like cells, with homozygous *PCDH19*-mutant derived neurons showing the longest neurites (**Fig. 5A**). These findings were in contrast with the molecular gene expression signatures which identified a set of genes that were differentially expressed only in the heterozygous *PCDH19*-mutant genotype. Gene expression in the brains of *Pcdh19*^mut/mut^ mice and homozygous *PCDH19*-mutant derived neurons were more similar to their respective WT genotype (**Figs 2,3,5**). It is possible that the homogeneous cellular environment of neurons derived from hESCs cannot fully replicate the brain’s complex cellular environment and for this reason the homozygous *PCDH19*-mutant derived neurons demonstrated differences in neurite outgrowths compared to WT cells. In the brain, multiple cell types interact and provide diverse physical and chemical cues, and these interactions are difficult to reproduce in vitro. As a result, neurite outgrowth observed in culture dishes may not accurately represent neurite development in the brain. Nonetheless, several of the genes exhibiting differential expressions only in the heterozygous phenotypes, were validated in the human cellular model even though the morphology of the *PCDH19*-mutant derived neurons was different. Importantly, some genes showed differential expression in both heterozygous and homozygous *PCDH19*-mutant derived neurons, such as the HOX gene family and other genes (**Supplementary Figs 3-4**).

RNA sequencing of brain samples from mice with different *Pcdh19* mutant genotypes showed differential expression in genes involved in sodium/potassium channel and activity, immediate early genes (IEGs), calcium mediated signalling, transmembrane function, NTRK signalling, cognitive function, and development (**Fig. 3A-D** and **Supplementary Fig. 3A**). Differential expression in several *Hox* genes was detected, including *Hoxa4, Hoxd3, Hoxa5, Hoxb5, Hoxb4* and *Hoxc5* (**Supplementary Fig. 3C**). The activation of *Hox* genes during differentiation was one of the highly enriched pathways in the analysis (**Fig. 3A)**. The expression of these genes was found to increase significantly in *Pcdh19*^WT/mut^ and *Pcdh19*^mut/mut^ mice, which is in accordance with the “heterochrony” or difference in timing hypothesis. According to this hypothesis, the *Pcdh19*^WT/mut^ condition can lead to asynchrony because of differences in speed of development between the WT expressing cells and the PCDH19 mutant expressing cells, which may lead to downstream abnormalities in neuronal networks that can manifest epileptic seizures. It was shown that loss of PCDH19 increases neurogenesis in mouse neural progenitor cells (mNPCs) and human neural progenitor cells (hNPCs)^7,26,35^. PCDH19-CE patient cells which were reprogrammed and differentiated into neurons showed advanced stage of maturation compared to their WT counterparts. In mosaic cultures, the development was classified as non-uniform/asynchronized while in WT cultures, it was classified as uniform/synchronized^26^. These results aligned with another study which showed increased neurogenesis in mNPCs through inhibition of *Pcdh19* expression in mouse cortices^26^. Thus, the altered functions of PCDH19 in early stages of development kinetics or trajectory of NPCs can contribute to the pathology in PCDH19-CE.

Another group of genes that were differentially expressed were genes involved in transmembrane transport, such as *Slc6a5, Slc14a2, Slc6a12, Slc18a3* and *Slc4a4*. Expression of these genes was significantly higher in *Pcdh19*^WT/mut^ mice brains compared to *Pcdh19*^WT/WT^ mice brains. However, the expression of *SLC18A3* was found to be downregulated in human heterozygous *PCDH19*-mutant derived neurons compared to WT and homozygous *PCDH19*-mutant derived neurons (**Supplementary Fig. 4**). This is in contrast to the increased expression of *Slc18A3* in the brains of *Pcdh19^WT/mut^* mice (**Fig. 3F**). These SLC transporters play a vital role in synaptic vesicle cycle and transport, neurotransmitter release, and transsynaptic signalling. Increased activity of these genes can lead to hyperactivity, ultimately causing seizures^36,37^. These results support the involvement of signalling pathways in manifesting the PCDH19-CE phenotype.

Genes in several additional pathways, related to synaptic transmission, myelination, sodium-calcium ion channel dependent activity, were found to be upregulated in the brains of *Pcdh19*^WT/mut^ mice. While genes in other pathways, such as voltage gated potassium channels, nerve growth factor (NGF)-stimulated transcription, NTRK signalling, NPAS4 mediated regulation of target genes, BDNF signalling and binding to members of canonical WNT pathway, were downregulated (**Figs 3, 5**). These findings are consistent with previous reports of disruption in synaptic transmission, post-presynaptic signalling and ion channel in primary culture of *Pcdh19*-knock out mouse hippocampal neurons^11,12,20,38^.

One interesting gene that was found to be significantly downregulated in both brains of *Pcdh19*^WT/mut^ mice and human heterozygous *PCDH19*-mutant derived neurons, is *Tmem40*. Not much is known about *Tmem40* except for its up-regulation and possible involvement in several types of cancer^39,40,41^. Here, for the first time, expression of *Tmem40* has been shown to be correlated with *PCDH19* mutation status and to be strongly down-regulated only in the heterozygous genotype in both mice and humans (**Figs 3E, 5C**).

It was recently reported that *PCDH19* mutation leads to enrichment in proteins involved in cytoskeleton organisation and actin binding^42^. We identified Myo3b, a gene which is involved in regulation of actin filament length and cytoskeleton organisation, to be downregulated in brains of *Pcdh19*^WT/mut^ mice (**Fig. 2D**). This downregulation might affect the binding of PCDH19 with β-catenin and disrupt WNT signalling pathway. Along with this, PCDH19 has also been shown to interact with regulators of RHO-GTPases^43^. We also identified several genes that were differentially expressed in brains of *Pcdh19*^WT/mut^ mice that are involved in RHO-GTPases binding, microtubule cytoskeleton disorganization, synaptic impairment and signalling pathways. It is very plausible that the defect in neurites length in the human heterozygous *PCDH19*-mutant derived neurons is related to impaired Rac/Rho signalling.

It has been previously described that GABA receptors play an important role in PCDH19-CE, but we did not find any differential expression of GABA genes in our RNA sequencing analysis^24,25^. However, genes of the pathways involved in synaptic transmission, and pre and post synaptic potential were significantly downregulated in brains of *Pcdh19*^WT/mut^ mice compared to *Pcdh19*^WT/WT^ and *Pcdh19*^mut/mut^ mice.

In this work we used a cellular human neuronal model system to both confirm our mouse results and to characterize the neuronal phenotype of the different *PCDH19* genotypes. In contrast to what was found in mutant *Pcdh19* mice brain samples, we found that expression of some IEGs, including *ARC, EGR2, NR4A1* and *C-FOS,* increased in heterozygous and homozygous human *PCDH19*-mutant derived neurons **(Supplementary Fig. 4)**. This is in accordance with a previous study that showed increased expression of IEGs on shRNA-mediated *Pcdh19* knockdown in mouse hippocampal neurons^11^. The increase in expression of IEGs in homozygous *PCDH19*-mutant derived neurons was observed for the first time. However, the expression of *NPAS4,* a known regulator of IEGs, was lower in both brains of *Pcdh19*^WT/mut^ mice and in human heterozygous *PCDH19*-mutant derived neurons compared to homozygous *PCDH19*-mutant clones and WT cells (**Figs 3 and 5C**). *NPAS4* expression was previously reported to increase in shRNA-mediated *Pcdh19* knockdown^11^. Using the human *PCDH19*-mutant neuronal like cells we observed that the neurite lengths of both heterozygous and homozygous *PCDH19*-mutant clones are significantly longer than WT clones (**Fig. 5A, B**).

Taken together, our RNA sequencing analysis of brains of *Pcdh19*-mutant mice and cellular studies of human cells suggests that PCDH19 is involved in signalling pathways and synaptic transmission and identified changes in the expression of several downstream genes which are unique to the heterozygous genotype. Future experiments that directly measure neuronal activity of the different genotypes will be highly informative for the correlation between genotype and phenotype. Potential targeted therapeutic experiments should focus on targeting enriched pathways such as upregulating the levels of *TMEM40* or other genes which are downregulated in the *PCDH19*^WT/mut^ genotype. Another therapeutic approach is to use *PCDH19* antisense oligonucleotides to downregulate *PCDH19* levels in heterozygous *PCDH19* mice/patient cells, with the aim of reversing the phenotype so it is more similar to the homozygous *PCDH19* null phenotype that resembles the WT.

## Acknowledgements

The authors wish to thank Hila Porat for inspiring us to work on the PCDH19-CE disorder and raising awareness and funding for part of this research. The authors also wish to acknowledge funding by Israel Precision Medicine Program (IPMP) ISF Research Grant No. 2039/20 to R.K.

## Author contributions

RK designed and supervised the study; EE and RJ designed, engineered and generated the mouse and human genetically modified ES cells, respectively. SK performed most of the cellular assays, the gene expression analysis and its validation. MA performed several cellular and gene expression validations. RK, SK, MA, and ZS wrote the manuscript.

## References

1. Chilcott E, Díaz JA, Bertram C, Berti M, Karda R. Genetic therapeutic advancements for Dravet Syndrome. Epilepsy and Behavior. 2022;132. doi:10.1016/j.yebeh.2022.108741

2. Ding J, Wang L, Jin Z, et al. Do All Roads Lead to Rome? Genes Causing Dravet Syndrome and Dravet Syndrome-Like Phenotypes. Front Neurol. 2022;13. doi:10.3389/fneur.2022.832380

3. Hoshina N, Johnson-Venkatesh EM, Hoshina M, Umemori H. Female-specific synaptic dysfunction and cognitive impairment in a mouse model of PCDH19 disorder. Science. 2021;372(6539). doi:10.1126/science.aaz3893

4. Smith L, Singhal N, El Achkar CM, et al. PCDH19-related epilepsy is associated with a broad neurodevelopmental spectrum. Epilepsia. 2018;59(3):679–689. doi:10.1111/epi.14003

5. Dibbens LM, Tarpey PS, Hynes K, et al. X-linked protocadherin 19 mutations cause female-limited epilepsy and cognitive impairment. Nat Genet. 2008;40(6):776–781. doi:10.1038/ng.149

6. Kolc KL, Møller RS, Sadleir LG, Scheffer IE, Kumar R, Gecz J. PCDH19 Pathogenic Variants in Males: Expanding the Phenotypic Spectrum. Adv Exp Med Biol. 2020;1298:177–187. doi:10.1007/5584_2020_574

7. Borghi R, Magliocca V, Petrini S, et al. Dissecting the Role of PCDH19 in Clustering Epilepsy by Exploiting Patient-Specific Models of Neurogenesis. J Clin Med. 2021;10(13). doi:10.3390/jcm10132754

8. Pederick DT, Homan CC, Jaehne EJ, et al. Pcdh19 Loss-of-Function Increases Neuronal Migration In Vitro but is Dispensable for Brain Development in Mice. Sci Rep. 2016;6. doi:10.1038/srep26765

9. Biswas S, Emond MR, Jontes JD. Protocadherin-19 and N-cadherin interact to control cell movements during anterior neurulation. J Cell Biol. 2010;191(5):1029–1041. doi:10.1083/jcb.201007008

10. Emond MR, Biswas S, Jontes JD. Protocadherin-19 is essential for early steps in brain morphogenesis. Dev Biol. 2009;334(1):72–83. doi:10.1016/j.ydbio.2009.07.008

11. Gerosa L, Mazzoleni S, Rusconi F, et al. The epilepsy-associated protein PCDH19 undergoes NMDA receptor-dependent proteolytic cleavage and regulates the expression of immediate-early genes. Cell Rep. 2022;39(8). doi:10.1016/j.celrep.2022.110857

12. Mincheva-Tasheva S, Nieto Guil AF, Homan CC, Gecz J, Thomas PQ. Disrupted Excitatory Synaptic Contacts and Altered Neuronal Network Activity Underpins the Neurological Phenotype in PCDH19-Clustering Epilepsy (PCDH19-CE). Mol Neurobiol. 2021;58(5):2005–2018. doi:10.1007/s12035-020-02242-4

13. Hayashi S, Inoue Y, Hattori S, et al. Loss of X-linked Protocadherin-19 differentially affects the behavior of heterozygous female and hemizygous male mice. Sci Rep. 2017;7(1). doi:10.1038/s41598-017-06374-x

14. Lamers D, Landi S, Mezzena R, et al. Perturbation of Cortical Excitability in a Conditional Model of PCDH19 Disorder. Cells. 2022;11(12). doi:10.3390/cells11121939

15. Giansante G, Mazzoleni S, Zippo AG, et al. Neuronal network activity and connectivity are impaired in a conditional knockout mouse model with PCDH19 mosaic expression. Mol Psychiatry. 2024;29(6):1710–1725. doi:10.1038/s41380-023-02022-1

16. Park J, Lee E, Kim CH, Ohk J, Jung H. Mosaicism-independent mechanisms contribute to Pcdh19-related epilepsy and repetitive behaviors in Xenopus. Proc Natl Acad Sci U S A. 2024;121(21). doi:10.1073/pnas.2321388121

17. Robens BK, Yang X, McGraw CM, et al. Mosaic and non-mosaic protocadherin 19 mutation leads to neuronal hyperexcitability in zebrafish. Neurobiol Dis. 2022;169. doi:10.1016/j.nbd.2022.105738

18. Cwetsch AW, Ziogas I, Narducci R, et al. A rat model of a focal mosaic expression of PCDH19 replicates human brain developmental abnormalities and behaviours. Brain Commun. 2022;4(3). doi:10.1093/braincomms/fcac091

19. Lv X, Ren SQ, Zhang XJ, et al. TBR2 coordinates neurogenesis expansion and precise microcircuit organization via Protocadherin 19 in the mammalian cortex. Nat Commun. 2019;10(1). doi:10.1038/s41467-019-11854-x

20. Giansante G, Mazzoleni S, Zippo AG, et al. Neuronal network activity and connectivity are impaired in a conditional knockout mouse model with PCDH19 mosaic expression. Mol Psychiatry. 2024;29(6):1710–1725. doi:10.1038/s41380-023-02022-1

21. Park J, Lee E, Kim CH, Ohk J, Jung H. Mosaicism-independent mechanisms contribute to Pcdh19-related epilepsy and repetitive behaviors in Xenopus. Proc Natl Acad Sci U S A. 2024;121(21). doi:10.1073/PNAS.2321388121

22. Robens BK, Yang X, McGraw CM, et al. Mosaic and non-mosaic protocadherin 19 mutation leads to neuronal hyperexcitability in zebrafish. Neurobiol Dis. 2022;169. doi:10.1016/J.NBD.2022.105738

23. Cwetsch AW, Ziogas I, Narducci R, et al. A rat model of a focal mosaic expression of PCDH19 replicates human brain developmental abnormalities and behaviours. Brain Commun. 2022;4(3). doi:10.1093/BRAINCOMMS/FCAC091

24. Serratto GM, Pizzi E, Murru L, et al. The Epilepsy-Related Protein PCDH19 Regulates Tonic Inhibition, GABAAR Kinetics, and the Intrinsic Excitability of Hippocampal Neurons. Mol Neurobiol. 2020;57(12):5336–5351. doi:10.1007/s12035-020-02099-7

25. Bassani S, Cwetsch AW, Gerosa L, et al. The female epilepsy protein PCDH19 is a new GABAAR-binding partner that regulates GABAergic transmission as well as migration and morphological maturation of hippocampal neurons. Hum Mol Genet. 2018;27(6):1027–1038. doi:10.1093/hmg/ddy019

26. Homan CC, Pederson S, To TH, et al. PCDH19 regulation of neural progenitor cell differentiation suggests asynchrony of neurogenesis as a mechanism contributing to PCDH19 Girls Clustering Epilepsy. Neurobiol Dis. 2018;116:106–119. doi:10.1016/j.nbd.2018.05.004

27. Pederick DT, Richards KL, Piltz SG, et al. Abnormal Cell Sorting Underlies the Unique X-Linked Inheritance of PCDH19 Epilepsy. Neuron. 2018;97(1):59–66.e5. doi:10.1016/j.neuron.2017.12.005

28. 28. Gecz J, Thomas PQ. Disentangling the paradox of the PCDH19 clustering epilepsy, a disorder of cellular mosaics. Curr Opin Genet Dev. 2020;65:169–175. doi:10.1016/j.gde.2020.06.012

29. Zhang Y, Pak CH, Han Y, et al. Rapid single-step induction of functional neurons from human pluripotent stem cells. Neuron. 2013;78(5):785–798. doi:10.1016/j.neuron.2013.05.029

30. 30. Kim D, Pertea G, Trapnell C, Pimentel H, Kelley R, Salzberg SL. TopHat2: accurate alignment of transcriptomes in the presence of insertions, deletions and gene fusions. Genome Biol. 2013;14(4). doi:10.1186/gb-2013-14-4-r36

31. Anders S, Pyl PT, Huber W. HTSeq-A Python framework to work with high-throughput sequencing data. Bioinformatics. 2015;31(2). doi:10.1093/bioinformatics/btu638

32. Love MI, Huber W, Anders S. Moderated estimation of fold change and dispersion for RNA-seq data with DESeq2. Genome Biol. 2014;15(12). doi:10.1186/s13059-014-0550-8

33. Zhang Y, Pak CH, Han Y, et al. Rapid Single-Step Induction of Functional Neurons from Human Pluripotent Stem Cells. Neuron. 2013;78(5):785–798. doi:10.1016/J.NEURON.2013.05.029

34. Dibbens LM, Tarpey PS, Hynes K, et al. X-linked protocadherin 19 mutations cause female-limited epilepsy and cognitive impairment. Published online 2008. doi:10.1038/ng.149

35. Mincheva-Tasheva S, Pfitzner C, Kumar R, et al. Mapping combinatorial expression of non-clustered protocadherins in the developing brain identifies novel PCDH19-mediated cell adhesion properties. Open Biol. 2024;14(4). doi:10.1098/rsob.230383

36. Aykaç A, Şehirli AÖ. The Role of the SLC Transporters Protein in the Neurodegenerative Disorders. Clin Psychopharmacol Neurosci. 2020;18(2):174–187. doi:10.9758/cpn.2020.18.2.174

37. Gil-Perotín S, Jaijo T, Verdú AG, et al. Epilepsy, status epilepticus, and hemiplegic migraine coexisting with a novel SLC4A4 mutation. Neurol Sci. 2021;42(9):3647–3654. doi:10.1007/s10072-020-04961-x

38. Lin Y, Bloodgood BL, Hauser JL, et al. Activity-dependent regulation of inhibitory synapse development by Npas4. Nature. 2008;455(7217):1198. doi:10.1038/nature07319

39. Zhang ZF, Zhang HR, Zhang QY, et al. High expression of TMEM40 is associated with the malignant behavior and tumorigenesis in bladder cancer. J Transl Med. 2018;16(1):9. doi:10.1186/s12967-017-1377-3

40. Liu D, Zhou G, Shi H, Chen B, Sun X, Zhang X. Downregulation of Transmembrane protein 40 by miR-138-5p Suppresses Cell Proliferation and Mobility in Clear Cell Renal Cell Carcinoma. Iran J Biotechnol. 2020;18(1):e2270. doi:10.30498/ijb.2019.85193

41. Zhang ZF, Liu F, Zhang HR, et al. Upregulation of TMEM40 is associated with the malignant behavior and promotes tumor progression in cervical cancer. Discover Oncology. 2023;14(1):43. doi:10.1007/s12672-023-00648-9

42. de Nys R, Gardner A, van Eyk C, et al. Proteomic analysis of the developing mammalian brain links PCDH19 to the Wnt/β-catenin signalling pathway. Mol Psychiatry. 2024;29(7):2199–2210. doi:10.1038/s41380-024-02482-z

43. Emond MR, Biswas S, Morrow ML, Jontes JD. Proximity-dependent Proteomics Reveals Extensive Interactions of Protocadherin-19 with Regulators of Rho GTPases and the Microtubule Cytoskeleton. Neuroscience. 2021;452:26–36. doi:10.1016/j.neuroscience.2020.09.033

