## Supplementary figures and images for "Molecular characterization of the heterozygous loss of function mutations in the X-linked PCDH19 gene causing PCDH19-Cluster Epilepsy"

### sup.1.jpg

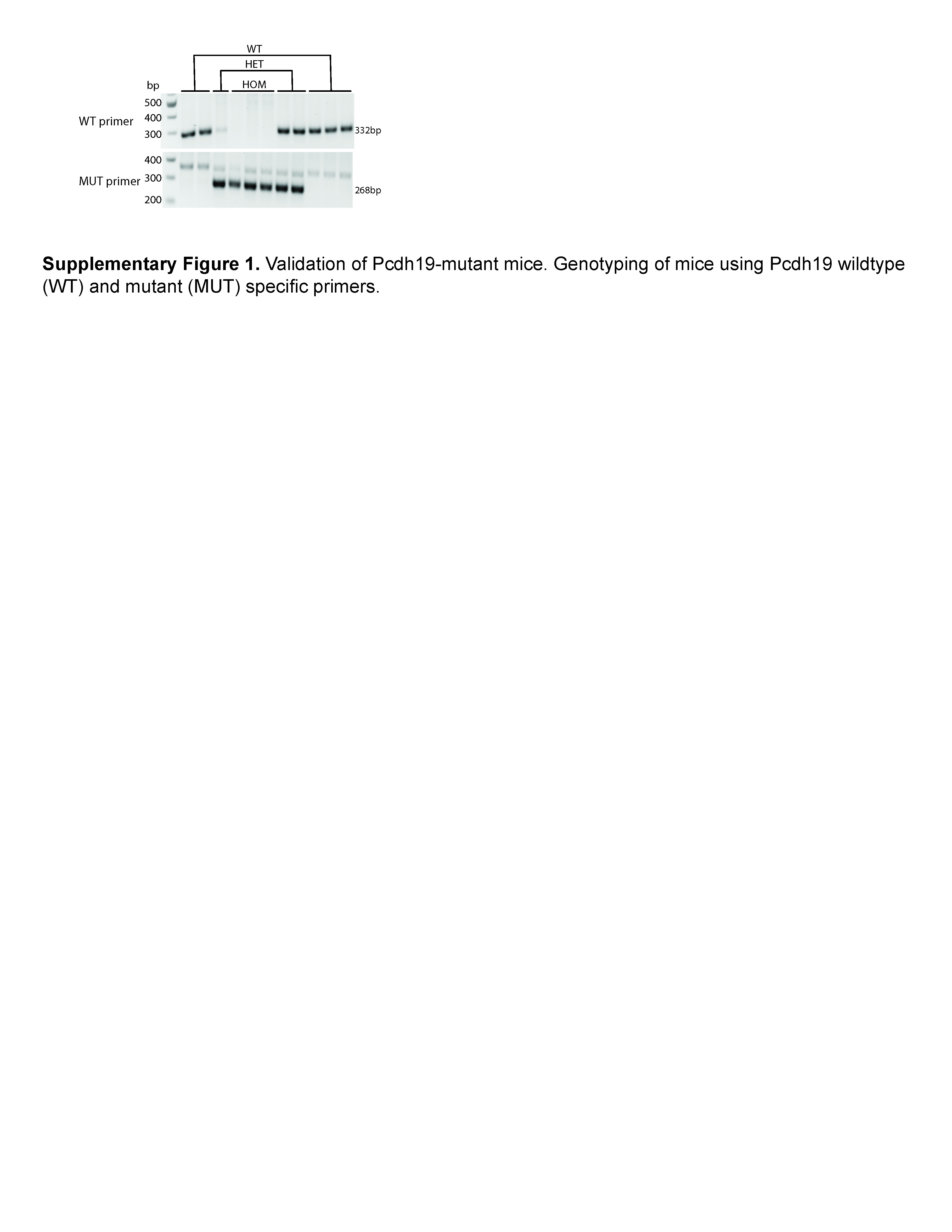

### Sup.2.jpg

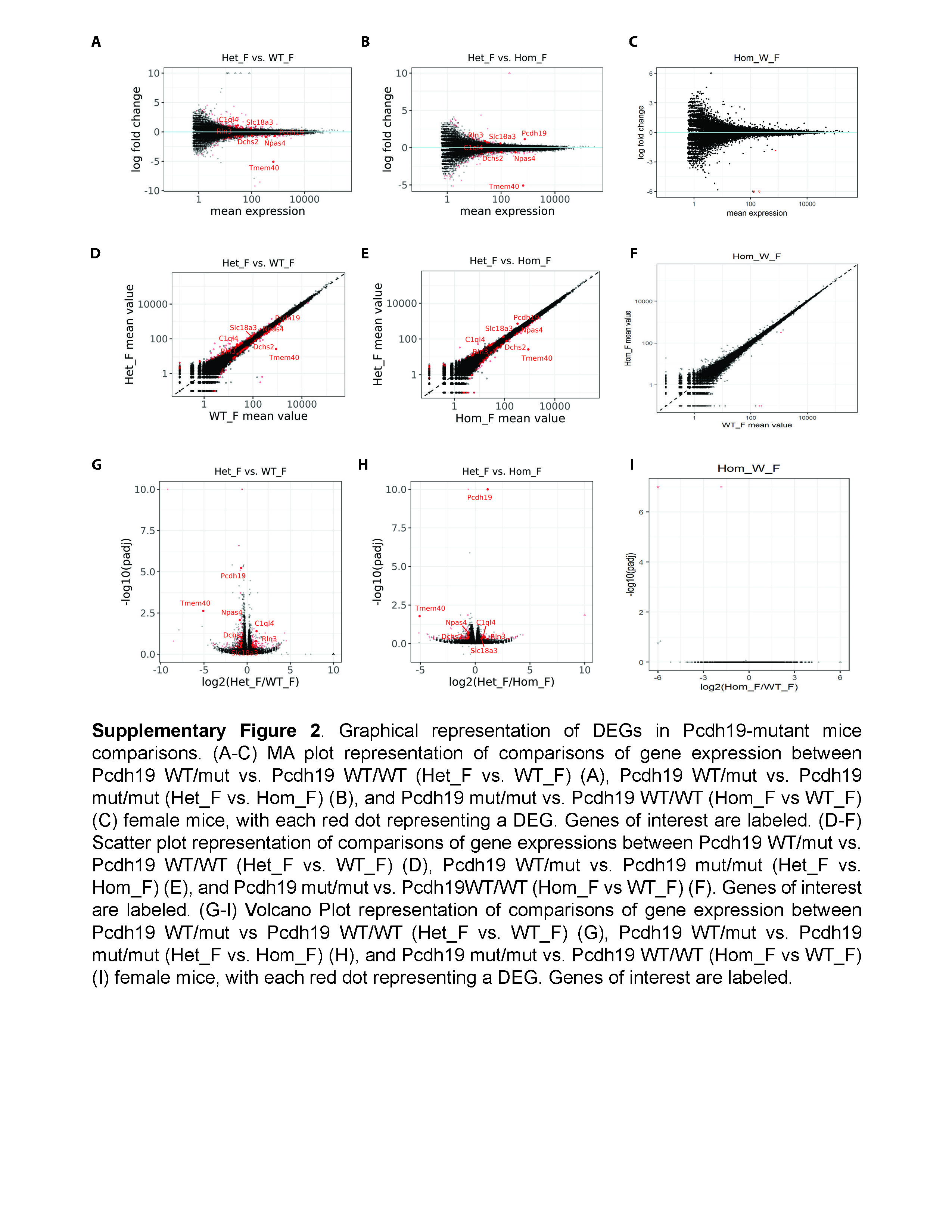

### Sup.3.jpg

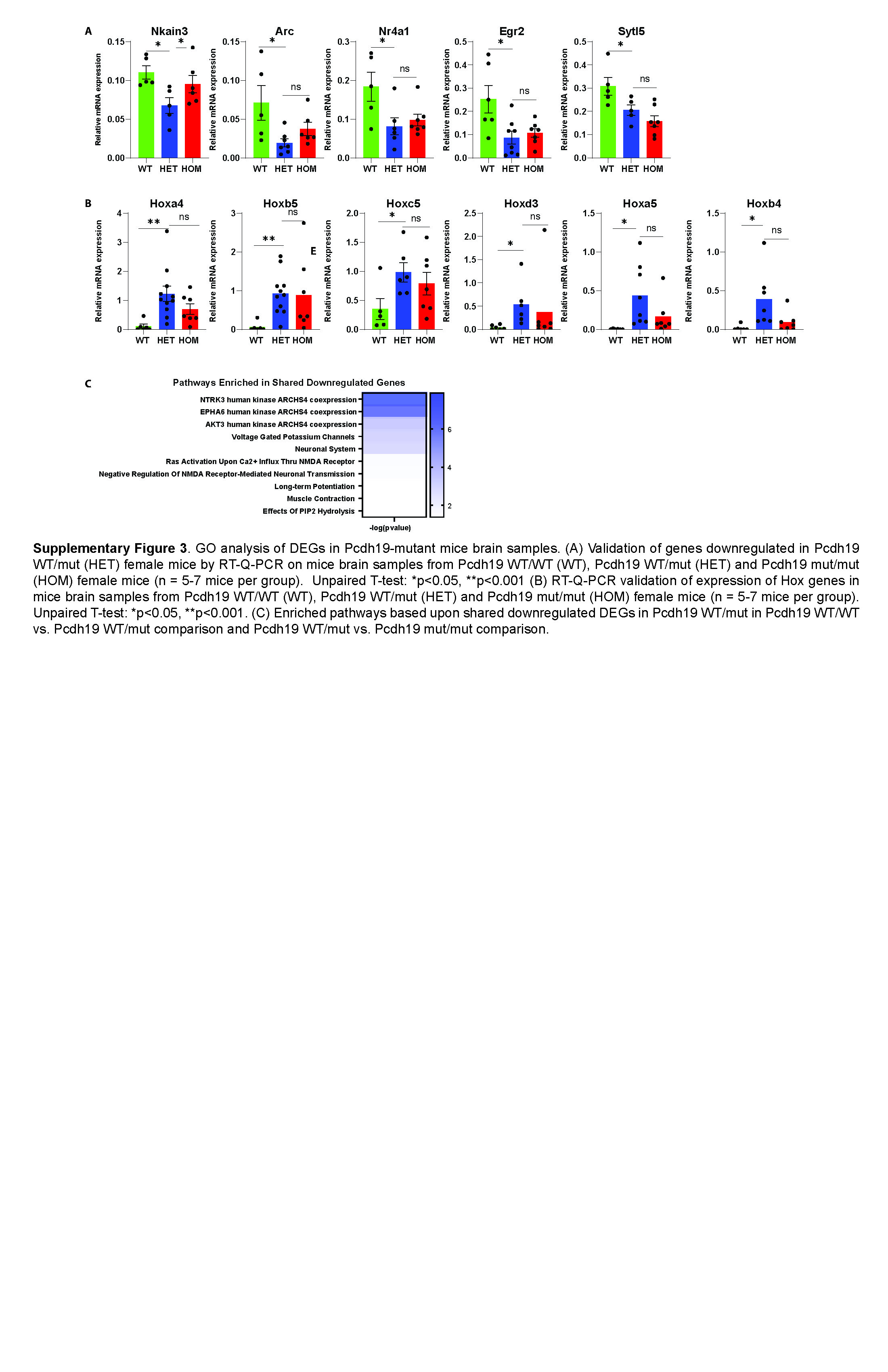

### Sup.4.jpg

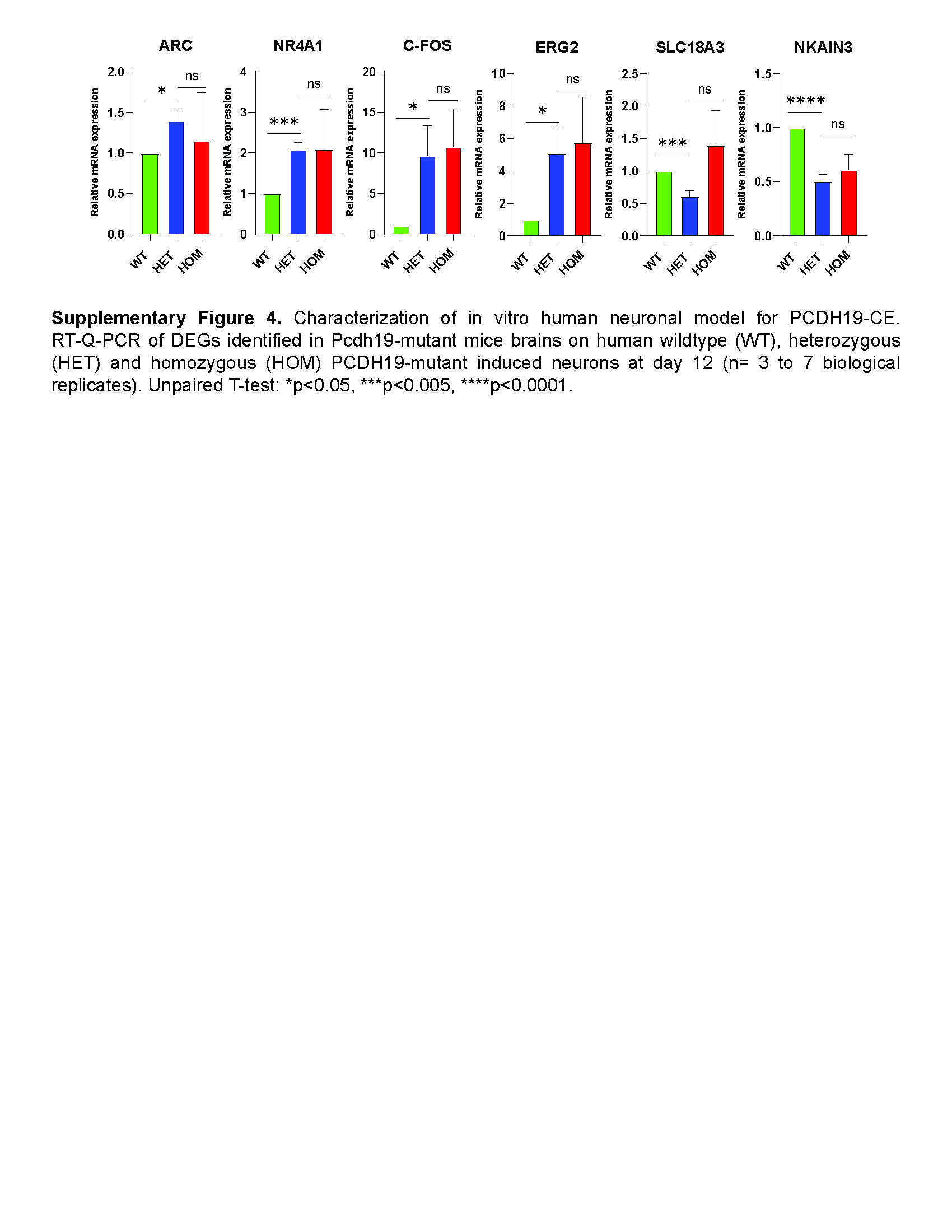
